# ExpressUrself: A spatial model for predicting recombinant expression from mRNA sequence

**DOI:** 10.1101/2022.12.02.518907

**Authors:** Michael P. Dunne, Javier Caceres-Delpiano

## Abstract

Maximising the yield of recombinantly expressed proteins is a critical part of any protein engineering pipeline. In most cases, the expression of a given protein can be tuned by adjusting its DNA coding sequence, however finding coding sequences that optimise expression is a nontrivial task. The 3-dimensional structure of mRNA is known to strongly influence the expression levels of proteins, due to its effect on the efficiency of ribosome attachment. While correlations between mRNA structure and expression are well established, no model to date has succeeded in effectively utilising this information to accurately predict expression levels. Here we present ExpressUrself, a model designed to capture spatial characteristics of the sequence surrounding the start codon of an mRNA transcript, and intended to be used for optimising protein expression. The model is trained and tested on a large data set of variant DNA sequences and is able to predict the expression of previously unseen transcripts to a high degree of accuracy.

## 1 Introduction

Proteins are macromolecules that are central to almost every aspect of cellular function, including cell structure, enzymatic reactions, cellular transport, signalling, and cell defence. Their ubiquity makes them not only key targets for study in a huge range of disciplines but also prime candidates for industrial production and usage in biotechnology, with applications ranging from medicines [1] to washing powders [2] and food ingredients [3].

The industrial manufacture of proteins relies on recombinant expression, i.e. the production of a protein of interest within a host organism that is easier to manipulate and faster to grow than the source organism, but which has not evolved to produce that protein at all. Each protein product requires specifically tailored production pipelines to ensure they are fast, safe, and cost effective [4]. In particular, tuning the expression rate of a given protein, i.e. the amount of protein produced, is of vital importance, and optimising it for maximal protein production is a far from trivial task [4].

The standard molecular biology dogma states that information flows from DNA into RNA (transcription) and finally into protein (translation). The expression levels of a recombinantly expressed protein can be affected by many interacting factors at all stages of this process, including: the choice of expression system [5, 6]; the structure, sequence, and properties of the protein being expressed [7]; the fitness of the host organism in the presence of the final protein product [8]; the choices of codons used in the RNA sequence [9]; and geometric properties of the transcribed mRNA [10, 11]. Processes for optimising these factors typically involve a considerable degree of trial and error [4, 12] over several iterative stages of expensive, time-consuming laboratory experiments.

For many practical purposes in protein engineering, the greatest degree of freedom, and therefore the primary target for optimising the expression of a given protein, is the choice of its underlying DNA.coding sequence [13]. In particular, it is the rate of translation that is typically tuned. Currently, the most common method of maximising translation rates is to use codon optimisation, which matches the sequence’s codon distribution to the distribution of tRNAs produced by the host organism [4]. However, codon optimisation is a complex, poorly understood, and often *ad hoc* process that does not reliably produce optimal DNA sequences [14, 15].

In contrast, it is well established that the 3D structure of RNA can have a significant impact on the expression levels of a protein, due to its impact on start codon accessibility [16, 17, 18, 19]. The contribution of mRNA structure to expression is not straightforward, and can be affected by multiple competing features near the start codon (Figure 1). Although correlations have been observed between expression and derived structural properties of the mRNA (such as minimum free energy), there has been limited progress in using either these or the sequence directly to reliably predict recombinant protein expression levels. The most relevant study to date, by Nikolados *et al* [20], which used the same data as is used in this work, was able to predict expression well for sequences similar to those seen in the training set, but did not generalise well to unseen sequences.

**Figure 1:**
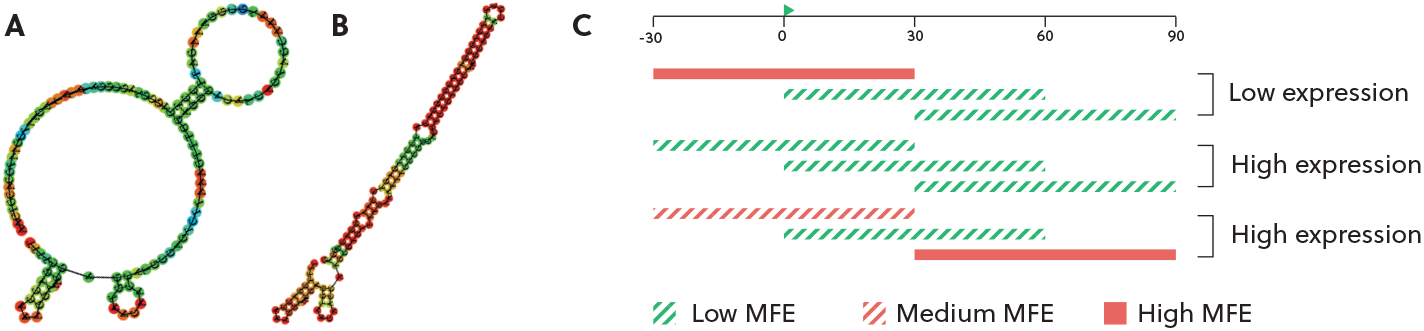
Minimum free energy (MFE) can be used as a measure of RNA structure strength. A, B) Examples of RNA structures for sequences with high and low MFEs respectively; C) Sequences that are highly structured around the start region are often poorly expressed, however this can be compensated by more tightly structured regions further downstream [19].

The model presented here, ExpressUrself, aims to predict the expression levels of a protein given only the mRNA at positions -30 to +90 relative to the start codon. It is trained on a large data set of GFP expression measurements [19], and consists of a sequence processing step adapted from UFold^1^ [21], followed by UNet, convolutional, and feed-forward layers. The ExpressUrself method is designed to be used comparatively between two sequences with the same amino acid sequence and expression system. In particular, it is designed to allow fast iterative optimisation of a protein’s DNA sequence to maximise the rate of translation initiation and thus increase levels of recombinant expression.

## 2 Data, model architecture, and training

### 2.1 Data description

The data set used to train the model was taken^2^ from a study by Cambray *et al*. [19], which measured the luminosity of 244,000 GFP constructs that had each had a different 96-nucleotide leader sequence attached after the start codon and before the coding region. The study was designed in a combinatorial fashion to evaluate several different candidate properties, including structural properties of three different regions between positions -30 and +90 relative to the start codon. This was the region that we chose as input to the model, and the luminosity value *p*_*NI*_ was used as the model output.

### 2.2 Train/test split

To prevent the model from recognising sequence-specific signals in the test set, the data was prepared such that sequence similarity between the train/validation and test sets was kept as low as possible. Firstly, each sequence in the source data is present in three replicates with minor variations of between 1 and 4 nucleotides, with a mean pairwise difference of 2.35 nucleotides. To avoid essentially triplicating each data point, we chose to use only the first replicate (rep1) of each sequence. Secondly, the data set is arranged into 56 mutational series, with each series containing several thousand variants of a single seed sequence. The sequences *within* mutational series exhibit high sequence similarity while pairs of sequences from different mutational series are always distinct. (Figure S1). The train, validation and test sets were chosen such that no sequence in any test set was drawn from the same mutational series as any sequence in the train or validation sets.

In order to achieve full coverage of the 56 mutational series while leaving unseen data for testing, the data set was randomly split into 14 different test groups, each containing four distinct series. In each case, models were trained on sequences from the other 52 series, and tested on the remaining 4 mutational series. The series used for each split are listed in Supplementary Table S1

### 2.3 Sequence preprocessing and model architecture

The input to the model is a 120-nucleotide sequence obtained by taking the 30 nucleotides preceding the start codon and the 90 nucleotides including and after it. The sequences were preprocessed using the same method as UFold [22]. This preprocessing step converts a sequence of length *ℓ* into a 17xLxL tensor, where L is the smallest multiple of 16 greater than *ℓ*. Here *ℓ* =120 and L=128. The preprocessing step involves taking a Kronecker product between a one-hot-encoded representation of the sequence and itself. The code was revised slightly to increase the speed of execution, from roughly 0.3s on a single CPU to 0.003s. This can be easily parallelised for multiple sequences.

The 17xLxL tensor is then inputted into a UNet module of height 4, which is inspired by the UFold method but reduced in size. We found that slightly reducing the size of the UNet module did not noticeably affect the performance of the model but increased the speed of training and inference considerably. The UNet layer is followed by convolutional layers and finally by a series of fully-connected layers. A diagram of the full architecture is shown in Figure 2.

**Figure 2:**
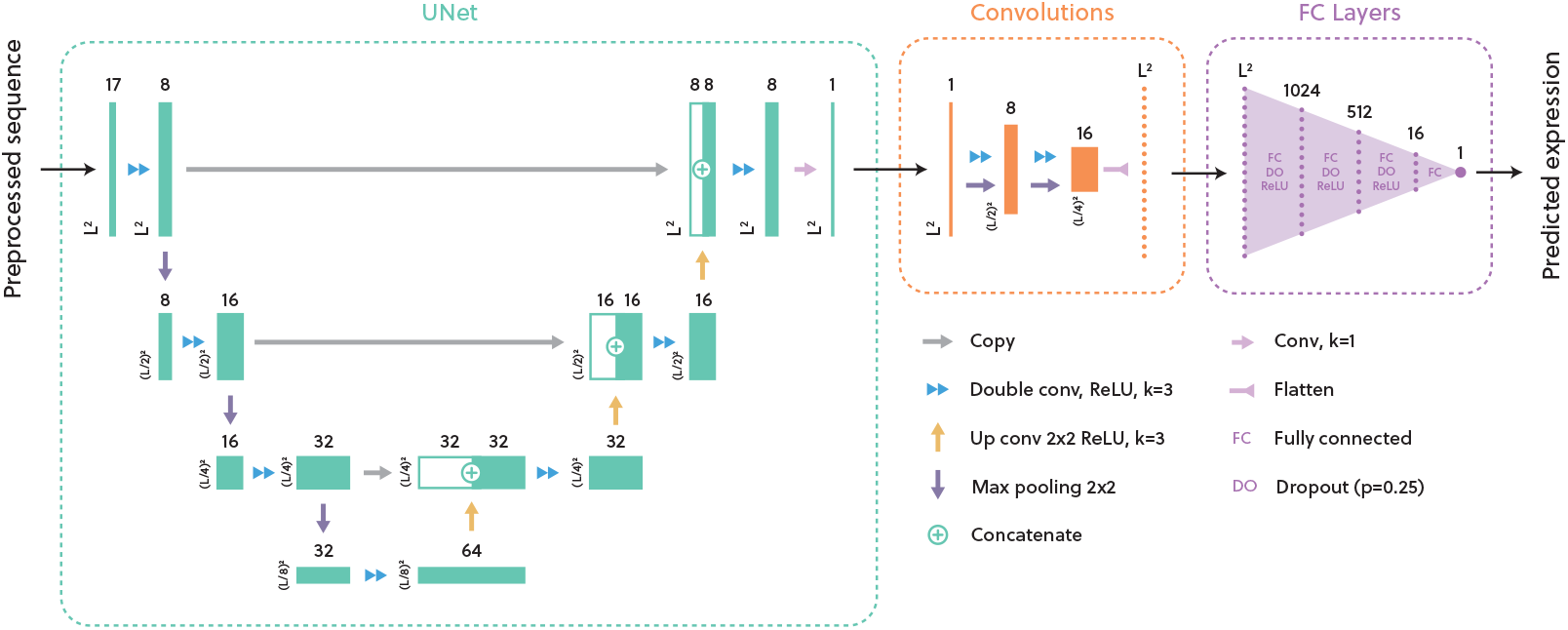
The model is formed of three separate blocks: a UNet module, followed by three double convolutional layers, followed by a series of fully connected layers. The input to the model is a preprocessed 17×128×128 tensor obtained from the mRNA sequence of the region -30:+90.

### 2.4 Data sampling and training

To reduce training time during development, the data was sampled to 10% of the full data size. The data was first filtered to retain only the first replicate, and then 24400 data points were sampled such that values were as evenly distributed as possible across 99 equally sized bins of expression values.

The model was trained using MSE loss for a maximum of 100 epochs, with an early stopping time of 20. The model was trained three times on each series split. The hyperparameters for the model, including the sizes of the linear layers, were chosen via a grid search over various runs of a single training split (k52_rx1), to maximise the test *R*^2^ score. The batch size was 20; the learning rate was 10^*−*3^, with a weight decay of 10^*−*5^. A dropout probability of 0.25 was used in the linear layers during training. The model was trained on a single AWS NVIDIA Tesla K80 GPU. The final production model is an ensemble of all 14 models from the different series splits, with the option to restrict to a smaller ensemble depending on the time and computing constraints of the user.

## 3 Results

### 3.1 ExpressUrself achieves high correlations when trained on 10% of the data set

The model was trained and tested three times for each of the 14 series subsets. The model achieved high correlation values in all cases, with mean Spearman *ρ*, Pearson *r* and *R*^2^ values of 0.779, 0.780 and 0.602 respectively (Figure 3A). For each series subset, the trained model with the highest *ρ* values on the test set was chosen as the best model, and taken forward for further analysis.

**Figure 3:**
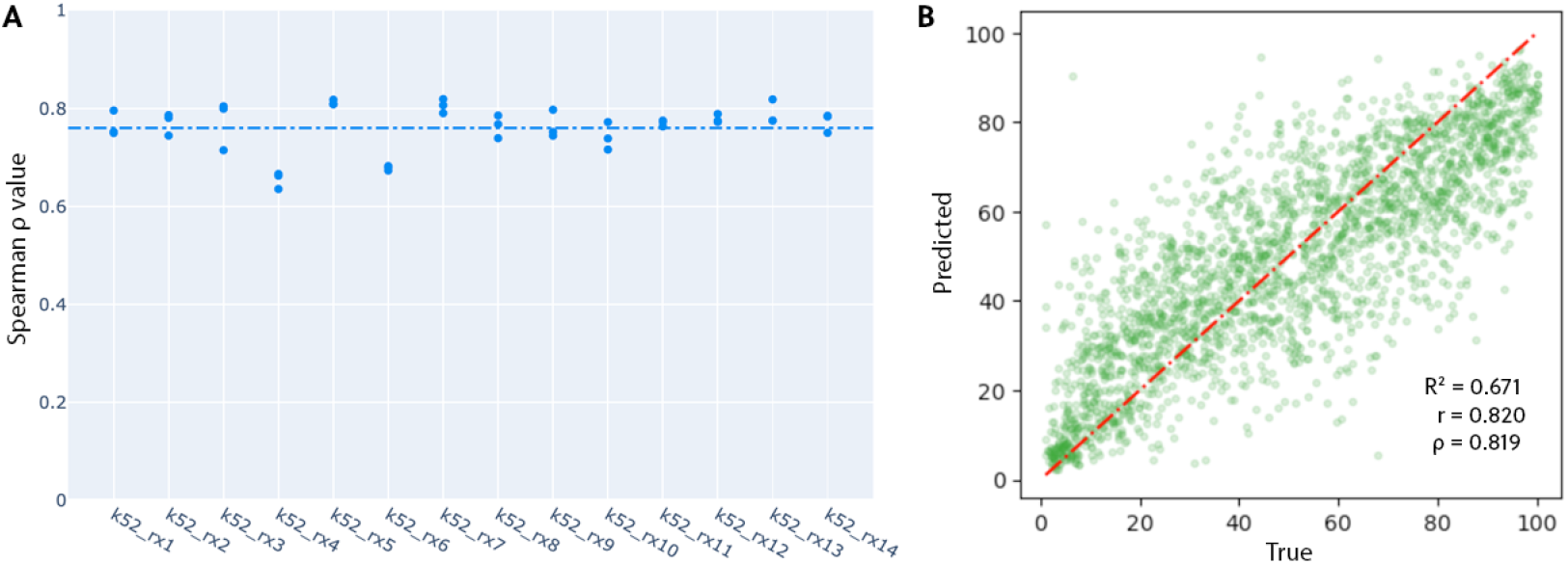
A) ExpressUrself has strong predictive power in each of the 14 series splits, with a mean Spearman *ρ* value of 0.779 (s.d. 0.046). The mean Pearson r and *R*^2^ values were 0.780 (s.d. 0.048) and 0.601 (s.d. 0.074) respectively. B) True vs. predicted values for predictions on series set k52_rx7. On this series, the model achieves *ρ*, r, and *R*^2^ values of 0.819, 0.820, and 0.671 respectively.

### 3.2 ExpressUrself categorises high- and low-expression sequences with high precision

On average across the best-performing models for each series split, ExpressUrself predictions that are very low (0 < y < 25) are correct (i.e. the true value is also in that range) in 78.9% of cases, compared with a baseline (defined as a model returning random values between 0 and 100) of 25% (see Figure 4A). Similarly, the model has a precision of 72.8% for very high (75 < y < 100) expression values. For y < 50 and y > 50, respectively, the model has a mean precision of 81.4% and 78.7%, compared with a baseline of 50%. This indicates that, even if there are some deviations in the values predicted by ExpressUrself, the broad classes of expression values predicted are likely to be correct.

**Figure 4:**
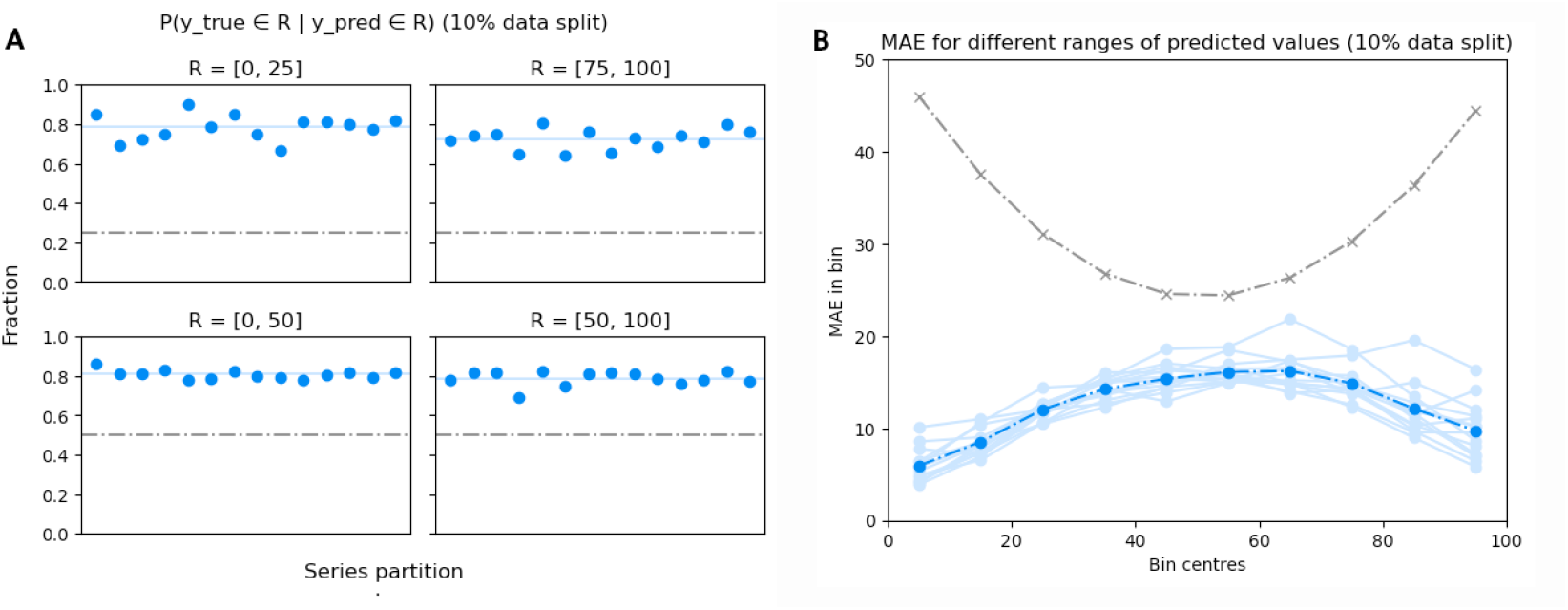
A) ExpressUrself identifies very low (y < 25), very high (y > 75), and low or high (y < 50 or y > 50) sequences with high mean precision values of 0.789, 0.728, 0.814, and 0.787 respectively.; B) The mean absolute errors are low compared to a random baseline (grey).

### 3.3 Low and high expression values are predicted with greater accuracy

In all of the series runs, ExpressUrself predictions that were further towards the edges of the data range were predicted with a lower mean absolute error (MAE): for predictions in the range [0, 10] the mean MAE across all series is 5.57; for [90, 100] it is 9.89, compared with 16.15 in the region [50, 60]. This is in contrast with what we would see if predictions were made at random (see Figure 4B, grey line), where edge values should be expected to have a much higher MAE.

## 4 Discussion

### 4.1 ExpressUrself excels at predicting extremal values

The ExpressUrself model has high precision and low MSE for low and high predicted values. It is likely that there exist some very strong signals for both low and high expression, such as particularly loose or tight RNA structures. In particular, tight, strong RNA structures covering the ribosome binding site are very unlikely to be expressed well. This would explain why the model seems to predict low values more accurately than high values. The results demonstrate that when extremal values are predicted by the model, they are likely to be in the correct region and should be trusted.

### 4.2 There is still a reasonably large margin of error for some sequences

The expression levels of a given gene depend on a huge range of factors, as well as being inherently dynamic. The data set used here was designed specifically to focus on sequence changes near to the start region, and also to keep larger-scale effects such as the protein sequence and structure and the expression system relatively constant. However, the model has been designed to pick up spatial characteristics of the RNA sequence, and so we must also assume that there are factors affecting expression that are contained in the sequence but which are simply inaccessible to the model.

For example, the model is unlikely to be able to pick up any effects relating to the elongation phase of translation, or relating to codon optimality. We saw no improvements in *R*^2^ when attempting to include codon or AT content into the models, however it is possible that the model could be improved by including these. Finally, there may be some sequences whose expression profiles are due to particularly complex interactions of multiple RNA structures either within the start region or with more distant parts of the mRNA transcript. Further work is required to tackle these tricky sequences.

### 4.3 ExpressUrself may struggle outside of the context of GFP expression

Although care was taken to ensure that the model was tested with sequences that were not used for training, its wider is currently unknown. Although it is not possible for the model to pick up sequence-specific motifs in the training data, it may be the case that the model is learning characteristics of the 5’ mRNA region that are specific to the expression system or experimental setup being used by Cambray *et al*., i.e. GFP expressed in *E. coli*. We believe the model is general enough to pick up universal characteristics of RNA and should be applicable to other experimental setups and organisms: we are currently working on testing and confirming this hypothesis in a laboratory.

## 5 Conclusion

Here we present ExpressUrself, a neural network model that is designed to extract spatial information relating to the structure of mRNA around the start codon of a transcript, and to use this to predict expression levels of the resultant protein. ExpressUrself has been trained on a large data set of variant GFP sequences, and tested on sequences that are completely distinct from the training set. Given that there are many factors that can affect the expression levels of a recombinantly expressed protein, the model is mostly intended for optimising gene sequences of a fixed protein in a fixed expression system. We are currently undertaking laboratory experiments to confirm ExpressUrself’s predictive power in settings outside of GFP constructs.

## Experimental data

We wish to thank Guillaume Cambray and his lab for generating the data used to train this model, as well as for helping us to interpret the data and for offering helpful suggestions.

## A Appendix

**Figure S1:**
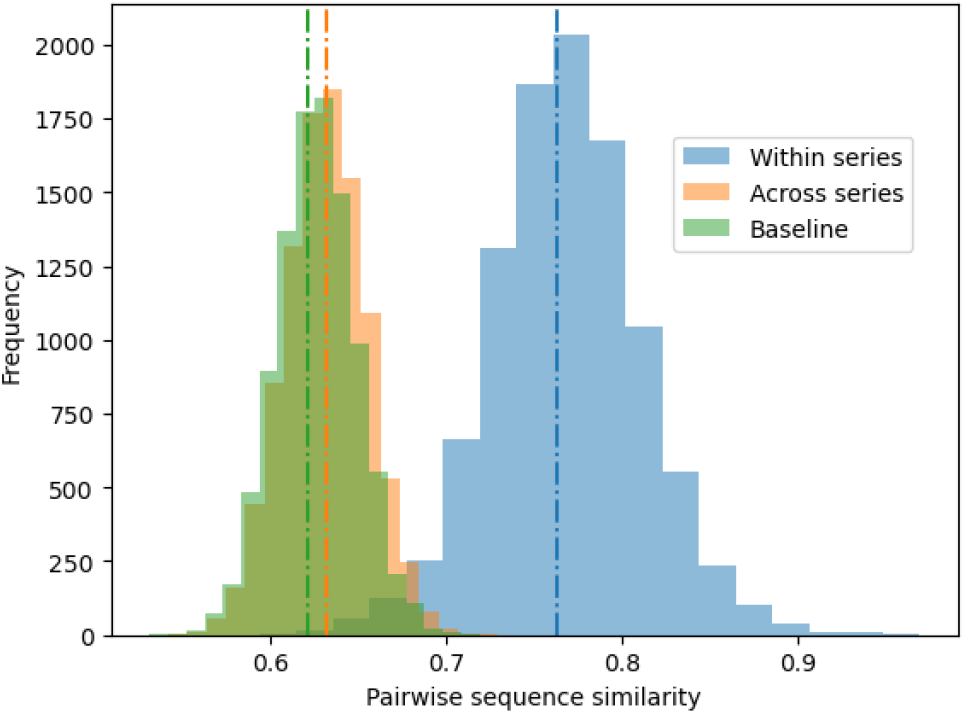
Distribution of sequence similarity scores for pairs of sequences from the same series (rep 1 only), across series, and, as a baseline, for pairs of randomly generated sequences. Sequence pairs were randomly sampled to a total of 10 000 pairs from each category.

**Table S1:** Series used for testing

| Subset ID | Test series | Subset ID | Test series |
| --- | --- | --- | --- |
| k52_rx1 | 44, 95, 133, 343 | k52_rx8 | 49, 54, 227, 314 |
| k52_rx2 | 20, 56, 86, 205 | k52_rx9 | 33, 84, 284, 340 |
| k52_rx3 | 1, 35, 76, 382 | k52_rx10 | 25, 26, 192, 359 |
| k52_rx4 | 15, 158, 344, 349 | k52_rx11 | 88, 117, 136, 379 |
| k52_rx5 | 50, 71, 99, 387 | k52_rx12 | 30, 102, 156, 355 |
| k52_rx6 | 60, 145, 221, 329 | k52_rx13 | 2, 47, 255, 384 |
| k52_rx7 | 247, 277, 299, 321 | k52_rx14 | 37, 46, 141, 380 |

## Footnotes

1 UFold is available under an MIT license at https://github.com/uci-cbcl/UFold

2 The data is available as a supplementary to the main paper by Cambray *et al*.

